# The variable ELF3 polyglutamine tract mediates complex epistatic interactions in Arabidopsis thaliana

**DOI:** 10.1101/061564

**Authors:** Maximilian Oliver Press, Christine Queitsch

## Abstract

Short tandem repeats are hypervariable genetic elements that occur frequently in coding regions. Their high mutation rate readily generates genetic variation contributing to adaptive evolution and human diseases. We recently proposed that short tandem repeats are likely to engage in epistasis because they are well-positioned to compensate for genetic variation arising at other loci due to their high mutation rate. We previously reported that natural ELF3 polyglutamine variants cause reciprocal genetic incompatibilities in two divergent *Arabidopsis thaliana* backgrounds. Here, we dissected the genetic architecture of this incompatibility and used a yeast two-hybrid strategy to identify proteins whose physical interactions with ELF3 were modulated by polyglutamine tract length. Using these two orthogonal approaches, we identify specific genetic interactions and physical mechanisms by which the *ELF3* polyglutamine tract may mediate the observed genetic incompatibilities. Our work elucidates how short tandem repeat variation, which is generally underascertained in population-scale sequencing, can contribute to phenotypic variation. Furthermore, our results support our proposal that highly variable STR loci can contribute disproportionately to the epistatic component of heritability.

## INTRODUCTION

Evolution is a tinkerer rather than a designer (Jacob 1977; Alon 2003); that is, adaptations are generally short-term, incremental fixes rather than alterations in fundamental biological plans. This principle is believed to underlie many design properties of biological systems. Thus, many (or most) genetic adaptations may be compensations for other genetic variants in a given background (Szamecz *et al.* 2014). One abundant source of genetic variation for such tinkering lies in short tandem repeats (STRs), genetic elements with high mutation rates. Due to these high mutation rates, STRs may be more likely than substitutions to contribute adaptive variants on a per-locus basis (Kashi *et al.* 1997; Gemayel *et al.* 2010; Hannan 2010). If ‘tinkering’ is a dominant mode of adaptation, STRs should be likely to show epistasis with other loci, and indeed, this expectation is borne out in the handful of well-characterized STRs (Press *et al.* 2014).

One such STR resides in the *Arabidopsis thaliana* gene *ELF3*, where it encodes a polyglutamine tract that varies in length across different natural strains (Tajima *et al.* 2007; Undurraga *et al.* 2012). We have previously shown that these *ELF3*-STR variants have strong effects on phenotype, and that these effects differ depending on the genetic background expressing a particular variant (Undurraga *et al.* 2012). These observations suggest that background-specific variants are modifying the effect of STR alleles through epistasis. The high variability of the *ELF3*-STR relative to expectations suggests that this STR may compensate for many background-specific polymorphisms across globally-distributed strains of *A. thaliana*.

ELF3 has been previously identified as a plausible candidate gene underlying a QTL for trait variance (*i.e.* noise) in the phenotypes of *A. thaliana* recombinant inbred lines (Jimenez-Gomez *et al.* 2011; Lachowiec *et al.* 2015). These results invite comparison to known ‘robustness genes’ such as HSP90 (Sangster *et al.* 2007, Sangster *et al.* 2008a; Sangster *et al.* b), which can reveal or conceal the phenotypic consequences of many other genetic variants. A mechanistic explanation of this robustness phenomenon is epistasis, in which a robustness gene interacts with many other loci (Queitsch *et al.* 2012; Lachowiec *et al.* 2015), as for the promiscuous chaperone HSP90 (Taipale *et al.* 2010). Our previous findings and the many studies describing ELF3’s crucial functions in plant development lead us to hypothesize that *ELF3* lies at the center of an epistatic network and that the ELF3’s polyglutamine tract modifies these interactions.

It is well-established that ELF3 functions promiscuously as an adaptor protein in multiple protein complexes that are involved in a variety of developmental pathways (Liu *et al.* 2001; Yu *et al.* 2008; Yoshida *et al.* 2009; Nusinow *et al.* 2011; Chow *et al.* 2012). Polyglutamine tracts such as the one encoded by the *ELF3*-STR often mediate protein interactions (Perutz *et al.* 1994; Stott *et al.* 1995; Schaefer *et al.* 2012). Therefore, it is plausible to assume that variation in the ELF3 polyglutamine tract affects ELF3’s interactions with its partner proteins. The ELF3 C-terminus, which contains the STR-encoded polyglutamine tract, is necessary for nuclear localization (Herrero *et al.* 2012) and ELF3 homodimerization (Liu *et al.* 2001), but thus far only one other protein (Phytochrome Interacting Factor 4, PIF4) has been shown to interact with this ELF3 domain (Nieto *et al.* 2014). Thus, the phenotypic and epistatic effects of ELF3-polyQ variation may arise from altered protein interactions, altered ELF3 nuclear localization, altered regulation of the PIF4 developmental integrator, or a combination thereof.

Here, we dissect the epistatic landscape modifying the function of the *ELF3*-STR through both physical and genetic interactions, and present evidence that this STR forms the hub of a complex network of epistasis, likely due to its role as a compensatory modifier of several other loci.

## METHODS

**Plant material and growth conditions:** Hypocotyl length was assayed in seedlings grown for 15d in incubators set to SD (8h light: 16h dark) at 22° on vertical plates as described previously (Undurraga *et al.* 2012). The *elf3-200* (Undurraga *et al.* 2012) and *elf3-4* (Hicks *et al.* 1996) mutants have been previously described. T-DNA lines (Alonso *et al.* 2003; Kleinboelting *et al.* 2012) were obtained from the Arabidopsis Biological Resource Center (Ohio State University).

<u>**Genotyping:**</u> For genotyping the *ELF3* STR and other loci across many F2 segregants, 1-2 true leaves from each seedling were subjected to DNA extraction. Seedlings were stored on their growth plates at 4° before genotyping but after phenotypic analysis. For genotyping the *ELF3* STR, PCR was performed in 10 *μ*L volume containing 0.5 *μ*M primers (Table S1), 0.2 *μ*M each dNTP, 1 *μ*L 10X ExTaq buffer, and 0.1 U ExTaq (Takara, Tokyo, Japan); with initial denaturation step of 95° for 5’, followed by 40 cycles of 95° 30”, 49° 20”, 72° 10”, with a final extension step at 72° for 5’. For other loci, PCR was performed in 20 *μ*L volume containing 0.5 *μ*M primers (Table S1), 0.2 *μ*M each dNTP, 2 *μ*L 10X ExTaq buffer, and 0.25 U Taq polymerase (NEB, Ipswich, MA); with initial denaturation step of 95° for 5’, followed by 35 cycles of 95° 30”, 55° 30”, 72° 1 ‘, with a final extension step at 72° for 5’.

**Genome resequencing:** Plants selected for genotyping-by-sequencing were transplanted to soil and grown under LD for 2-3 weeks. They were then stored at 4° until DNA extraction was performed. One late rosette-stage Ws individual was used for Ws whole-genome resequencing. DNA extraction was performed using the DNeasy Plant Mini kit (Qiagen, Valencia, CA) according to the kit protocol. This DNA was quantified using high-sensitivity Qubit fluorescence analysis (ThermoFisher Scientific, Waltham, MA) and re-genotyped with *ELF3*-STR primers (Table S1). We used 10 ng DNA from each F_2_ segregant in NextEra transposase library preparations (Illumina, San Diego, CA), or a standard 50 ng preparation for the Ws library. Library quality was assessed on a BioAnalyzer (Agilent, Santa Clara, CA) or agarose gels. The Ws individual was sequenced in one 300-cycle MiSeq v2 run (300 bp single-end reads) to ~12X coverage. The F_2_ segregant libraries were pooled and sequenced in one 200-cycle HiSeq v3 run to ~2X average coverage (100 bp paired-end reads, Table S3).

**Sequence analysis:** Reads were aligned to the Col reference genome using BWA v0.7.5 MEM (Li 2013), and variants were called using SAMtools v0.1.19 (Li *et al.* 2009). High-quality Ws variants (Q>=40) were thus identified from Ws parent data, and compared with variants in previously-sequenced related strains (Gan *et al.* 2011). F_2_ segregant genotype calls were combined into a single variant call format (VCF) file and filtered for loci with such Ws variants. We used SNPtools (Wang *et al.* 2013) to perform haplotype and genotype imputation for each locus in F_2_ segregants. For workflows employed in sequence analysis, see Supplementary Text. Following sequence analysis, one individual was found to be a heterozygote at the *ELF3* locus. This individual was omitted from all following analyses requiring *ELF3* homozygotes.

**Quantitative trait locus (QTL) analysis:** F_2_ genotypes were reduced to a set of 500 loci randomly sampled from the imputed genotypes, plus a single nucleotide variant (SNV) marking the *ELF3* locus. We used these genotypes to estimate a genetic map and perform QTL analysis using the R/qtl package (Broman *et al.* 2003). A nonparametric epistasis test was implemented in a custom R script using R/qtl functions. For a more detailed description of commands and the epistasis test, see the Supplementary Text. Follow-up genotyping of 10 additional F_2_s was performed using PCR markers (Table S1), and these genotypes were included in final QTL analyses.

**Candidate gene analysis:** Homozygous T-DNA lines (Alonso *et al.* 2003; Kleinboelting *et al.* 2012) with insertions in genes of interest were obtained from the Arabidopsis Biological Resource Center (Ohio State University) and phenotyped for hypocotyl length under SD at 15 days. All such experiments were performed at least twice. Double mutants were obtained by crossing relevant lines and genotyping (primers in Table S1). Mutant lines are listed in Table S2. Expression analysis confirmed no detectable *LSH9* expression in the *lsh9* mutant, suggesting that it is a null mutant. *LSH9* promoter sequences across strains were downloaded from the Salk 1,001 Genomes Browser (http://signal.salk.edu/atg1001/3.0/gebrowser.php) (Alonso-Blanco *et al.* 2016) and aligned using Clustal Omega v1.0.3 (Sievers *et al.* 2011).

**Yeast two-hybrid (Y2H):** *ELF3* variants with different STR lengths were PCR cloned out of cDNAs of previously described *A. thaliana* carrying *ELF3* transgenes (Undurraga *et al.* 2012) into the XmaI/BamHI sites of pGBKT7. Genes to be tested for ELF3 interactions were PCR cloned into the EcoRI/XhoI sites of pGADT7 from cDNAs of indicated strains (Table S1 for primers). Clones were confirmed by restriction digest and sequencing. The Y2H screen was performed against the Arabidopsis Mate and Plate cDNA library (Clontech, Madison, WI), essentially according to the manufacturer’s instructions, except selections were performed on C-leu-trp-his plates incubated at 23°. Clones which also showed activation of the *ADE2* reporter gene and did not autoactivate were subsequently tested against the various ELF3-polyQ constructs (see Supplementary T ext for details, full details on clones given in File S1).

LacZ activity was assayed through X-gal cleavage essentially as previously described (Möckli and Auerbach 2004), again in strains using PJ69-4a as Mata parent. For weakly activating constructs (GLDP1 and ELF4), 0.2 absorbance units of yeast were used in each assay to reduce background, and color development was assessed at points between 16 and 72 hours of incubation at room temperature.

**Quantitative PCR (qPCR):** For measuring *ELF3* and *LSH9* transcript levels, pooled aerial tissue of ~30 mg short-day-grown seedlings of each relevant genotype were collected at ZT8 7d post germination. RNA preparation and qPCR was performed as described previously (Undurraga *et al.* 2012), using primers in Table S1.

**Statistical analysis:** All statistical analyses and plotting was performed using R 2.15.3 or R 3.2.1 (R Core Team 2016). Analysis scripts are provided with data as detailed below.

**Data availability:** High-throughput sequencing data are available in BAM format at NCBI Sequence Read Archive accession SRP077615. Processed genotype data, phenotype data, and analysis code are available at https://figshare.com/s/e01a40b98a4ef5a9e5b3.

## RESULTS

Genetic analysis of *ELF3*-STR effects on hypocotyl length: To investigate the genetic architecture of epistasis for the *ELF3*-STR, we crossed two *A. thaliana* strains with a previously reported mutual incompatibility of their respective *ELF3*-STR variants (Col-0, Ws, (Undurraga *et al.* 2012)). We phenotyped the resulting F_2_ population for hypocotyl length under short days, a trait dramatically affected by ELF3 function. The Col and Ws backgrounds did not substantially differ in this trait (p = 0.16, Kolmogorov-Smirnov test, Figure 1A). Although most F_2_ seedlings showed phenotypes within the range of the two parental lines, the F_2_ phenotypic distribution showed a long upper tail of transgressive variation, and consequently a different distribution from either parent (p = 0.0039 against Ws, p = 0.055 against Col, Kolmogorov-Smirnov tests). As longer hypocotyls in light conditions indicate ELF3 dysfunction in the circadian clock (Liu *et al.* 2001), this observation is consistent with the co-segregation of incompatible Col and Ws alleles. We replicated this observation in a much larger population (1106 seedlings), which was used for further genetic analysis.

To investigate the genetic basis of the phenotypic transgression in hypocotyl length, we harvested the 720 most phenotypically extreme seedlings (longest and shortest hypocotyls) for genotyping (Figure S1). Each individual seedling was genotyped at the *ELF3* locus, using primers directly ascertaining the 27bp *ELF3*-STR-length polymorphism between Col and Ws. Across these individuals, we observed a strong main effect of the *ELF3* locus on phenotype (Figure 1B]), in which the Col allele of *ELF3* frequently showed transgressive phenotypes, though some individuals homozygous for the Ws allele also showed transgressive phenotypes. Specifically, a naÄve regression analysis of the data in Figure 1B indicated that each *ELF3-Col* allele increased hypocotyl length by 0.87±0.077 mm, and that the *ELF3* locus thereby explained 15% of phenotypic variation. This analysis is misleading, because it implies that Col seedlings should show longer hypocotyls than Ws seedlings due to *ELF3* genotype - this is not the case (Figure 1A).

Among these seedlings, the individuals with extreme phenotypes and individuals homozygous at the *ELF3* locus are expected to be most informative about **ELF3*-STR* effects on phenotype. Furthermore, *ELF3* genetic interactions are expected to be most apparent in *ELF3* homozygotes. Consequently, we used a novel genetic approach to detect epistasis between *ELF3* and other loci as follows. For each *ELF3* STR allele, we selected 24 homozygotes (Ws/Ws and Col/Col) at each phenotypic extreme (the shortest and longest hypocotyls). The sampling of extremes is an effective and statistically justified method for genetic mapping (Lander and Botstein 1989). These 96 individuals were analyzed in a genotyping-by-sequencing approach (Table S3, Figure S1). For details of this approach, see the Supplementary Information (Supplementary Text, Figures S2-S3).

**Figure 1.**
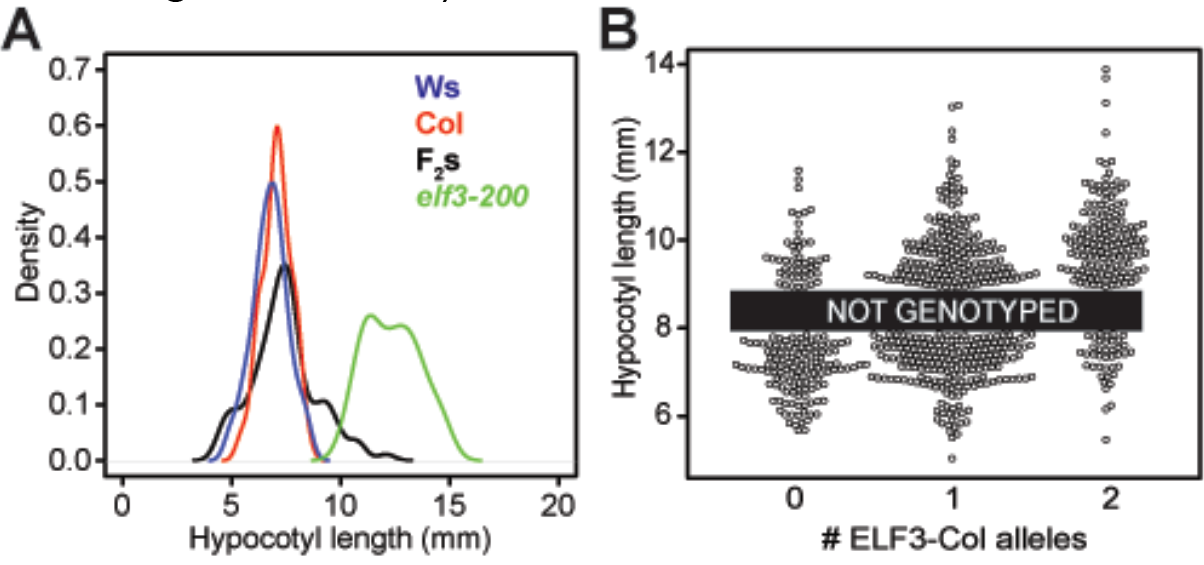
Phenotypic transgression in Col x Ws F_2_ segregants. (A): F_2_ segregant phenotypes compared to parents and an *elf3* null mutant in the Col background; n *≈* 50 for each homozygous line and n *μ* 100 for F_2_s. Colors indicate genotypes. Hypocotyl length was determined at 15d under short days. (B): Phenotypic distributions of a large population of Col x Ws F_2_ segregants, for 720 extreme individuals genotyped at the *ELF3* locus. N total = 1106 seedlings. 386 seedlings were not genotyped (indicated by the black box).

With these data, we performed a one-dimensional QTL scan to identify chromosomal regions contributing to hypocotyl length (Figure 2A). This analysis indicated a QTL on Chr2 corresponding to *ELF3* as expected, but also significant QTL on Chr1, Chr4, Chr5, and potentially one or more additional QTL on Chr2 affecting the phenotype. A two-dimensional QTL scan suggested that at least some of these QTL interact epistatically with the *ELF3* locus (Figure S4).

We binned F_2_s homozygous at *ELF3* according to their *ELF3* genotype, and performed one-dimensional QTL scans on each homozygote group separately (masking the genotypes of all other individuals). We observed that the same LOD peaks were replicated well in *ELF3*-Col homozygotes, but poorly in *ELF3-*Ws homozygotes (Figure 2B). Notably, a second Chr2 QTL was thus revealed, indicating that loci other than *ELF3* on Chr2 are relevant to the phenotype (at least in *ELF3*-Col plants). This analysis suggested that the *ELF3*-STR genotype is epistatic with at least four other loci controlling this phenotype, with effects masked in *ELF3*-Ws plants.

**Figure 2.**
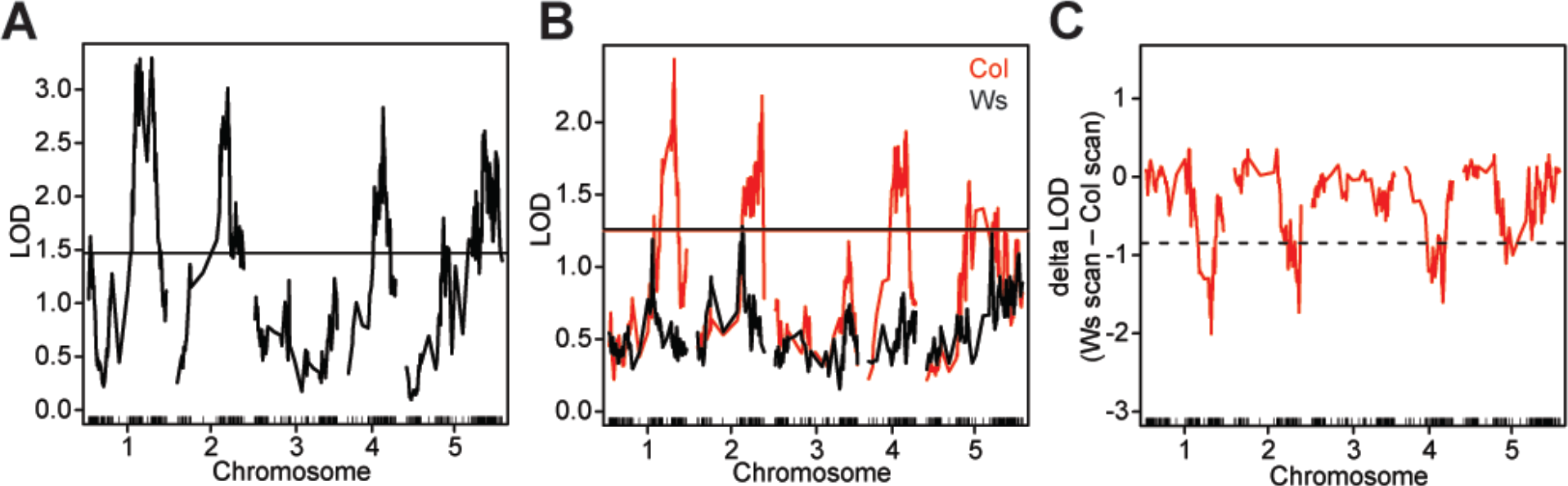
QTL analysis identifies interactions of *ELF3* with multiple loci. (A): onedimensional QTL scan including all sequenced F_2_s. Horizontal line indicates 99% significance threshold based on permutations. (B): QTL scan stratified by *ELF3*-STR genotype (all genotypes but those of indicated F_2_s masked in each analysis). Horizontal lines of each color indicate 99% significance threshold based on permutations for each scan. (C): A nonparametric test of epistasis between *ELF3* and other loci, using the independent QTL scans shown in (B). *ELF3* is located on chromosome 2. Dotted horizontal line indicates 99% significance threshold based on permutations.

To directly test for epistasis with *ELF3*, we adapted a previously described method (Sangster *et al.* 2008a). Separating the *ELF3* homozygotes again, we used permutations to define an empirical null distribution for the difference of likelihood (LOD) scores expected between the Ws and the Col scans. When comparing the difference in LOD scores between the two QTL scans at all loci, we found that the peaks on Chr1, Chr2, and Chr4 (and to a lesser extent Chr5) were all stronger in Col (Figure 2C). Consequently, these loci constitute background-specific *ELF3* interactors.

We considered the genetic contribution of these loci to the phenotype using a multiple QTL mapping approach, using both the independently estimated QTL locations and a refined model re-estimating QTL positions based on information from all QTLs (Table S4). In each case, loci of strong effect on Chr1, Chr2, and Chr4 were supported, along with interactions between Chr2 (*ELF3*) and the other two loci. In the refined model, the Chr5 locus and the second (other than *ELF3*) Chr2 locus were also strongly supported. We conclude that although *ELF3* interacts epistatically with a variety of other loci in determining hypocotyl length, the principal contributors to *ELF3*-mediated effects on the trait are on Chr1 and Chr4. Moreover, direct inspection of phenotypic effects of *ELF3* in interaction with each putative locus among F_2_ segregants supported the hypothesis of epistasis with *ELF3* most clearly for the Chr1 and Chr4 loci (Figure S5).

**Candidate gene analysis identifies *LSH9* as a genetic interactor of *ELF3:*** The chromosome intervals identified by our QTL analysis encompassed a large number of genes, and overlapped with an inversion between these backgrounds on Chr4 (Rowan *et al.* 2015). Previous work using a multiparent *A. thaliana* mapping population (Kover *et al.* 2009) also identified possible candidate genes affecting hypocotyl length in these regions (Khattak 2014).

We phenotyped mutants of several candidate genes in the Col background under the conditions of our intercross experiment (15d SD hypocotyl length, Figure S6). We observed small phenotypic effects of the T-DNA insertion mutants *lsh9* and *nup98*. However, these small effects on their own cannot explain the transgressive phenotypic variation in F_2_s (Figure 1A).

**Figure 3.**
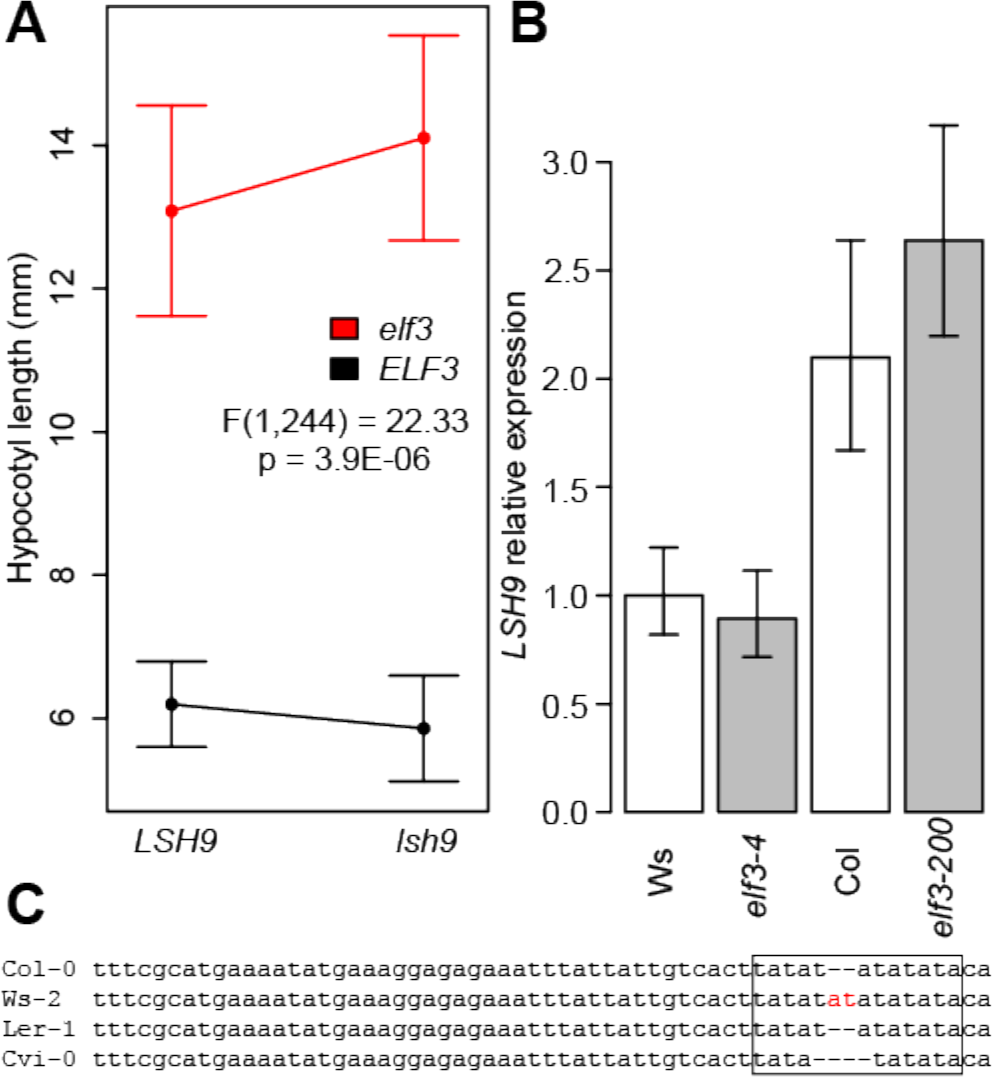
*ELF3* interacts genetically with *LSH9*, which shows background-specific expression. (A): Double mutant analysis of *elf3* and *lsh9* seedlings grown for 15d under SD. ANOVA analysis of the interaction between the mutant effects on phenotype is displayed. Error bars indicate standard deviation, n > 35 for each genotype. (B): qRT-PCR analysis of *LSH9* transcript levels across genotypes. Seedlings were grown under SD and harvested at ZT8 (dusk). *LSH9* expression is expressed as a proportion of Ws expression, normalized relative to *UBC21*, error bars are standard error from three technical replicates. Experiments were repeated with similar results. (C): Ws-specific STR polymorphism in the *LSH9* promoter. The -221 to -163 region (relative to start codon) of the *LSH9* promoter is aligned across diverse *A. thaliana* strains (Col, Ws-2, Ler-1, Cvi-0), with the small [TA]_n_ STR boxed. Ws-2 is a separately maintained stock of the Ws (Wassilewskija) strain. The Ws-2-specific polymorphism is highlighted in red.

We generated double mutants between these mutants and the *elf3* null mutant to determine whether these genes interacted epistatically with *ELF3*. We found little evidence for an interaction between *nup98* and *elf3* mutations (Figure S6C). However, we detected a significant interaction between *ELF3* and *LSH9*, in the form of reciprocal sign epistasis between the two null mutants affecting hypocotyl length (Figure 3A). Although *lsh9* single mutants had significantly shorter hypocotyls than WT, *lsh9 elf3* double mutant hypocotyls were substantially longer than in *elf3* single mutants. *LSH9 (LIGHT-DEPENDENT SHORT HYPOCOTYLS 9)* is an uncharacterized gene belonging to a gene family named for LSH1, which is known to act in hypocotyl elongation (Zhao *et al.* 2004). Like other genes in this family, *LSH9* encodes a putative nuclear localization sequence but no other distinguishing features.

To test our hypothesis that ELF3-STR mediated epistasis may be due to altered protein interactions, we investigated whether LSH9 and ELF3 interacted physically using Y2H. However, we were unable to detect a physical interaction between the Col or Ws variants of LSH9 and ELF3 (Figure S7), suggesting a different mechanistic basis for the observed genetic interaction. For example, *LSH9* expression may depend on ELF3 function as a transcriptional regulator. Alternatively, *ELF3* expression may depend on LSH9 function. We tested both hypotheses by measuring expression levels of *LSH9/ELF3* in, respectively, *elf3* or *lsh9* mutant backgrounds (*elf3* mutants were available in both Col and Ws backgrounds, *lsh9* only in Col). *ELF3* expression levels were unchanged in *lsh9* mutants (Figure S8). Moreover, levels of *LSH9* transcript did not significantly differ between WT and *elf3* mutants in either strain background. However, *LSH9* expression was reduced in the Ws background relative to Col independently of *ELF3* genotype(Figure 3B). This result is consistent with the observed phenotypic interaction in F_2_s, which showed elongated hypocotyls when Col alleles at the *ELF3* locus co-segregated with Ws alleles at the *LSH9* locus (Figure S5), thereby pairing poorly-functioning *ELF3* alleles with potentially lower *LSH9* expression levels.

Taken together, *ELF3-LSH9* epistasis between Col and Ws may be due to regulatory changes between these two backgrounds altering *LSH9* transcript levels. Coincidentally, we observed that the *LSH9* promoter contains an STR polymorphism in the Ws background that may alter *LSH9* expression; alternatively *LSH9* altered expression in Ws may be due to trans-effects (Figure 3C).

**Figure 4.**
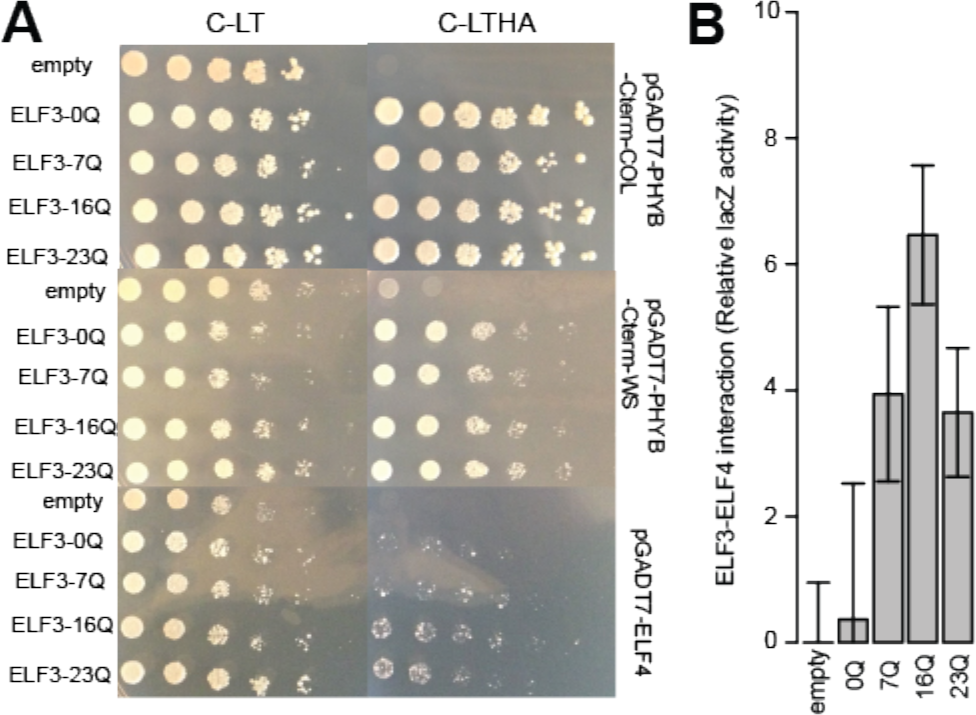
Y2H interaction of ELF3 with known protein interactors can be modulated by polyQ variation. (A): Yeast carrying indicated vectors were spotted in five-fold dilutions onto C-leu-trp (C-LT) or C-leu-trp-his-ade (C-LTHA) media. PHYB-Cterm: previously-defined C-terminal truncations of PHYB sufficient for ELF3 interaction (Liu *et al.* 2001) from the Col and Ws backgrounds. For each protein X, experiments were repeated with independent PJ69-4α + pGADT7-X transformants with similar results. (B): LacZ assays support polyQ effects on ELF3-ELF4 interaction. The strains shown in (A) also express *LacZ* from the Y2H promoter, whose activity was assayed in cell lysates (see Methods). In each assay, all observations are expressed relative to the activity of the empty vector, whose mean is set to 0. Error bars indicate standard deviation across three technical replicates. This experiment was repeated with similar results.

**ELF3-polyQ tract variation affects known protein interactions:** In parallel with our genetic analysis, we used Y2H to directly identify *A. thaliana* proteins whose physical interactions with ELF3 are polyQ-modulated. We first explored whether synthetic ELF3s with 0Q (no polyQ), 7Q (variant in Col), 16Q (variant in Ws), and 23Q (endogenous to strains Br-0 and Bur-0) forms of ELF3 show Y2H interactions with well-described ELF3 interactors PHYB (Liu *et al.* 2001), ELF4 (Nusinow *et al.* 2011; Herrero *et al.* 2012) (Figure 4), and PIF4 (Nieto *et al.* 2014). None of the ELF3 constructs showed autoactivation in yeast when paired with an empty vector (Figure S9). The ELF3-interacting domain of PHYB has two coding variants between Col and Ws, and we thus tested both Ws and Col variants of this domain. We found that both forms showed apparently equal affinity with all polyQ variants of ELF3. ELF4, which has no coding variants between Col and Ws, also interacted with all polyglutamine variants of ELF3, though rather weakly compared to PHYB. Under these conditions, a subtle preference of ELF4 for longer polyQ variants (e.g. ELF3-16Q and ELF3-23Q) was apparent. We confirmed this preference in a quantitative, growth-independent assay in which LacZ expression is driven by the Y2H interaction (Figure 4B).

We were not able to replicate the previously reported ELF3-PIF4 interaction (Nieto *et al.* 2014) for any ELF3-polyQ variant in our Y2H system (Figure S10), and were thus unable to evaluate effects of polyQ variation on ELF3-PIF4 interactions.

Together, our data suggest that ELF3-polyQ tract variation can affect ELF3 protein interactions, in particular if these interactions are weaker (as for ELF4) and presumably more sensitive to structural variation in ELF3.

**Figure 5.**
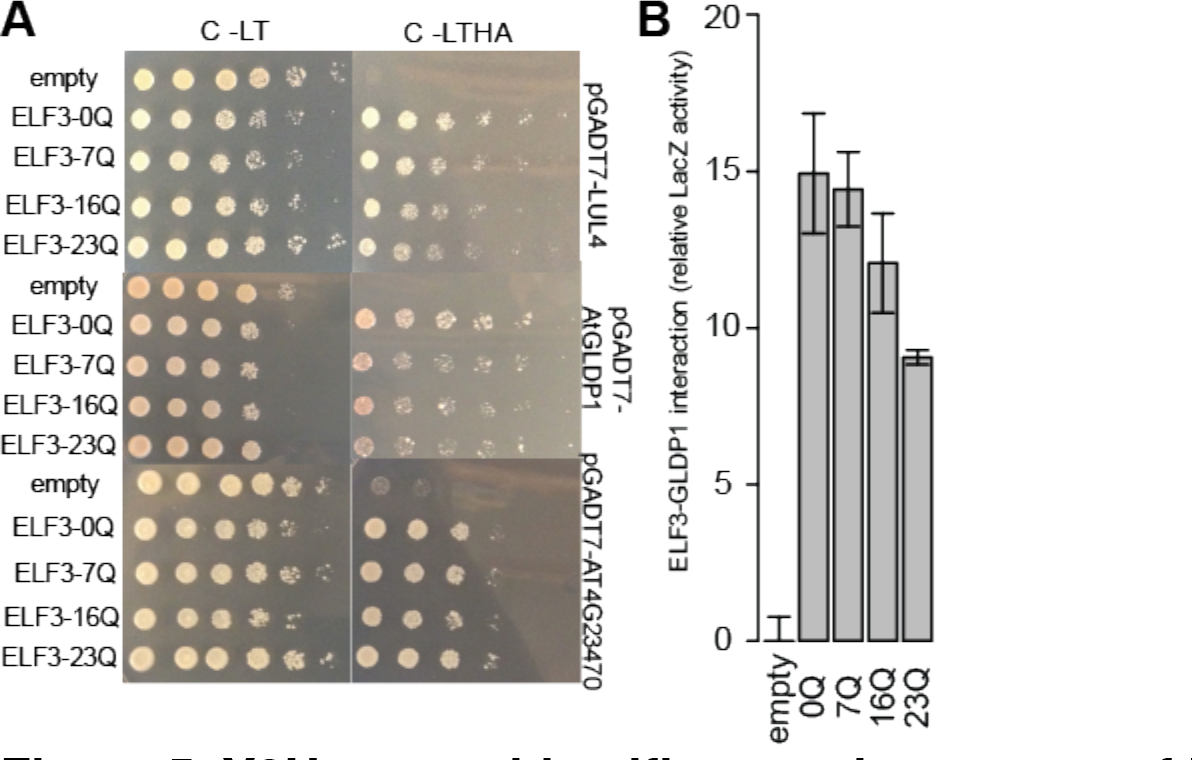
Y2H screen identifies new interactors of ELF3. (A): Yeast carrying indicated vectors were spotted in five-fold dilutions onto C-leu-trp (C-LT) or C-leu-trp-his-ade (C-LTHA) media. For each protein X, experiments were repeated with at least two independent PJ69-4α + pGADT7-X transformants with similar results. (B): LacZ assays support polyQ effects on ELF3-At-GLDP1 interaction. The strains shown in (A) also express *LacZ* from the Y2H promoter, whose activity was assayed in cell lysates (see Methods). In each assay, all observations are expressed relative to the activity of the empty vector, whose mean is set to 0. Error bars indicate standard deviation across three technical replicates. This experiment was repeated with similar results.

**Y2H screen identifies three novel ELF3 interactors, one of which is polyQ-modulated:** None of the known ELF3 interactors were encoded by genes located in the major Chr1 and Chr4 QTLs identified by our genetic screen. If the ELF3-polyQ tract mediates protein interactions, these regions should contain additional, previously-undescribed polyQ-modulated ELF3 interactors. We screened the ELF3-7Q protein for interactions with proteins from a commercially available library derived from Col, to detect ELF3-protein interactions within the Col background.

We subjected Y2H positives to several rounds of confirmation (Supporting Text), yielding a total of three novel proteins that robustly interacted with ELF3: PLAC8-domain-containing protein AT4G23470, LUL4, and AtGLDP1 (Figure 5). AT4G23470 was recovered in two independent clones, and LUL4 was recovered in three independent clones. The PLAC8-domain protein AT4G23470 is encoded by a gene within the QTL interval on chromosome 4, but this protein showed no variation in affinity among the various ELF3-polyQs. LUL4, a putative ubiquitin ligase, is not encoded in any of the mapped QTL and also shows no variation in affinity among the various ELF3-polyQs.Thus, differential interaction with these proteins is unlikely to underlie the observed epistasis.

In contrast, the AtGLDP1 protein, which is encoded on chromosome 4 but not within the QTL interval, appeared to show a subtle preference for the synthetic ELF3-0Q construct over longer polyQs tracts. We confirmed this preference in a quantitative LacZ assay (Figure 5B). Although our screen is unlikely to exhaust hitherto-unknown ELF3 interactors, our data suggest that the ELF3-polyQ tract can affect ELF3’s interactions with other proteins. Moreover, polyQ variation appears to affect weaker ELF3-protein interactions; strong protein interactions (for example ELF3-LUL4, ELF3-PHYB) are robust to polyQ variation.

## DISCUSSION

The contribution of STR variation to complex traits is thought to be considerable (Kashi *et al.* 1997; Press *et al.* 2014). Specifically, it has been proposed that STR variation contributes disproportionately to the epistatic term of genetic variance, due to its potential to contribute compensatory mutations. However, the molecular mechanisms by which different STRs contribute to genetic variance should derive from their particular features. For instance, polyQ variation may be expected to affect protein interactions (Perutz *et al.* 1994; Schaefer *et al.* 2012) or the transactivation activity of affected proteins (Escher *et al.* 2000). In this study, we considered the case of the previously-described *ELF3* STR (Undurraga *et al.* 2012).

We found that the genetic architecture of ELF3-dependent phenotypes is highly epistatic between the divergent Col and Ws strains, leading to substantial phenotypic transgression in the well-studied hypocotyl length trait. We identified at least 3 QTLs showing genetic interactions with the *ELF3* STR in a Col x Ws cross. These QTLs generally did not coincide with obvious candidate genes known to affect ELF3 function. Our confirmation of one genetic interaction (*LSH9*) in the Col background suggests that these QTLs encompass variants affecting hypocotyl length in tandem with *ELF3* STR variation. We cannot formally exclude the hypothesis that variants linked to *ELF3* (other than the Col and Ws *ELF3*-STR variants) may contribute to the observed phenotypic variation. For example, the ELF3-A362V-substitution in the *A. thaliana* strain Sha affects ELF3 function in the circadian clock (this site is invariant between Col and Ws) (Anwer *et al.* 2014). However, our previous work demonstrated that *ELF3*-STR variation suffices to produce strong phenotypic incompatibility between the Col and Ws background (Undurraga *et al.* 2012). Therefore, we reason that *ELF3*-STR variation is the most parsimonious explanation for the phenotypic variation, and in particular the observed transgression in the Col x Ws cross.

We further used Y2H screening to explore whether ELF3 polyQ tract variation affects protein interactions. ELF3’s promiscuous physical associations with other proteins are essential to its many functions in plant development (Liu *et al.* 2001; Kolmos *et al.* 2011; Nusinow *et al.* 2011; Herrero *et al.* 2012). Disruption of these interactions suggested an attractive mechanism by which ELF3 polyQ tract variation might affect ELF3 function. Assaying several known and novel ELF3-interacting proteins yielded evidence for a modest effect of polyQ variation on weaker protein interactions. However, there was no generic requirement for specific ELF3 polyQ tract lengths across all interactors. Indeed, the modest effects that we found were interactor-specific and thus not likely to generalize.

We did find that the ELF3-ELF4 interaction, which is crucial for circadian function and thus hypocotyl length (Nusinow *et al.* 2011; Herrero *et al.* 2012), demonstrates a subtle preference for the Ws 16Q ELF3 variant. However, there is no sequence variation in ELF4 between Col and Ws and we did not detect the ELF4 locus by QTL analysis, suggesting that this binding preference does not explain the transgressive phenotypes revealed by *ELF3*-STR variation (Figure 1A). However, the subtle polyQ-dependence of the ELF3-ELF4 interaction may play a role through indirect interactions.

Alternatively, rather than modulating ELF3 function as an encoded polyQ tract, the *ELF3*-STR may affect ELF3 transcription or processing. A previous study of an intronic STR in *A. thaliana* demonstrated that certain hyperexpanded STR alleles led to dysregulation of the IIL1 gene (Sureshkumar *et al.* 2009), presumably due to aberrant processing of IIL1 transcripts. Others have previously argued that such ‘informational’ (as opposed to ‘operational’) processes are more likely to have genetic or physical interactions (Jain *et al.* 1999). We have not tested this hypothesis; however, our previous studies found no correlation between *ELF3*-STR variation and *ELF3* expression across many natural strains (Undurraga *et al.* 2012).

Taken together, our findings support a model in which highly variable STRs can contribute to the epistatic component of heritability through both direct and indirect functional interactions with other loci.

**ELF3 is a model for sub-expansion polyQ variation:** PolyQ variation is best-known from hyperexpansions that are associated with several incurable human neurological disorders (Orr and Zoghbi 2007; Fondon *et al.* 2008; Usdin 2008; Hannan 2010), in which CAG (though generally not CAA) repeats dramatically elongate to lengths >50 units, reaching over 200 units in some patients. We argue that these hyperexpansion disorders are poor models for the functional impact of sub-expansion variation of polyQs tracts, and propose instead that the ELF3 polyQ might serve as a more appropriate model.

Although ELF3 polyQ tract variation can reach a length associated with disease in humans, it differs qualitatively from the well-studied human polyQ tract hyperexpansions. Human polyQ hyperexpansions are associated with protein aggregation or plaque formation, a phenomenon that requires a sufficiently long uninterrupted polyglutamine domain (Sharma *et al.* 1999; Lu and Murphy 2015). These previous *in vitro* studies suggest that although the ELF3 polyQ variants in Col and Ws are sufficiently different to alter phenotype, neither is long enough to lead to aggregation by these mechanisms (though the ELF3-23Q variant may be within this range).

Next, the effects of the disease-associated polyQ hyperexpansions are generally dominant, due to the nature of their molecular effects, which are generally thought to show a protein gain-of-function (Orr and Zoghbi 2007; Fondon *et al.* 2008). We observed no evidence that ELF3-polyQ variation behaves in a dominant fashion, but rather that Col x Ws F_1_s show approximately WT phenotypes. The effects of ELF3-polyQ variation manifest only when separated from a favorably interacting genetic context (as in segregating F_2_s).

Last, the (sometimes implicit) expectation from polyQ hyperexpansion phenotypes is that there is a linear, or at least monotonic, association between phenotypes and polyQ length. For instance, the degree of huntingtin polyQ expansion is strongly correlated with Huntington’s disease severity (ANDREW et al. 1993). In contrast, no ELF3-polyQ-related phenotype has been shown to have a monotonic association to ELF3 polyQ length (Undurraga *et al.* 2012; Press *et al.* 2014, Press *et al.* 2016). Indeed, all indications are that the mapping between ELF3 polyQ tract length and phenotype is non-monotonic and strongly contingent on genetic background, unlike the classic polyQ disease models. Our results suggest that sub-expansion polyQ tract variation can engage in multiple genetic interactions, and at least in some cases modulate protein interactions. More work is needed to evaluate the generality of our findings and determine the breadth of molecular mechanisms by which modest, sub-expansion polyQ variation can affect phenotype.

**ELF3 as a robustness gene**: Here, we operated under the assumption that a few strong polyQ-modulated interactions with ELF3 explain the polyQ-dependent genetic architectures. Alternatively, the ELF3 polyQ tract may modulate many transient interactions that are perturbed by hypomorphic ELF3 activity. In this interpretation, *ELF3* acts as a ‘robustness gene’ (Lempe *et al.* 2013). The best-described example of such is the protein chaperone HSP90 (Rutherford and Lindquist 1998), whose multiple transient interactions with many proteins-about 10% of the yeast proteome (Zhao *et al.* 2005)- lead to pleiotropic effects upon HSP90 inhibition or dysregulation (Sangster *et al.* 2007). ELF3 has been previously proposed as a robustness gene (Jimenez-Gomez *et al.* 2011), consistent with its promiscuity in protein complexes and the pleiotropic nature of *elf3* mutant phenotypes. Our finding that functional modulation of ELF3 by polyQ variation reveals several genetic interactors is consistent with this interpretation.

A similar hypothesis is that ELF3 “gates” robustness effects from robustness genes with which it interacts. For instance, we have recently shown that ELF3 function is epistatic to some of HSP90’s pleiotropic phenotypic effects (unpublished data, M. Zisong, P. Rival, M. Press, C. Queitsch, S. Davis), and ELF4 has also been proposed as a robustness gene governing circadian rhythms and flowering (Lempe *et al.* 2013). Here, we show that polyQ variation affects ELF3-ELF4 binding, which would provide a mechanistic link between ELF3 polyQ effects and a known robustness gene.

These hypotheses remain speculative in the absence of more explicit tests. Nonetheless, we suggest that the pleiotropic effects of polyQ variation in ELF3 (or similar cases) may be better understood by considering *ELF3* as a robustness gene, in which phenotypic effects are determined by a variety of important but individually small interactions of this highly connected epistatic hub.

## ACKNOWLEDGMENTS

We thank Karla Schultz and Katie Uckele for technical assistance. We thank Choli Lee and Jay Shendure for assistance with high-throughput sequencing of the Col x Ws F_2_ population and members of the Shendure laboratory for advice regarding library preparation. We thank Amy Lanctot for generating the pGBK-ELF3-0Q and pGBK-ELF3-23Q constructs. We thank Daniel Melamed and Stanley Fields for guidance in carrying out Y2H experiments and the generous gift of yeast strains. We thank Giang Ong and Maitreya Dunham for access to the MiSeq instrument for resequencing the Ws genome. We thank Stanley Fields and Evan Eichler for access to LightCycler instruments. We thank Elhanan Borenstein and members of the Queitsch and Borenstein laboratories for helpful conversations. MOP was supported in part by National Human Genome Research Institute Interdisciplinary Training in Genome Sciences Grant 2T32HG35-16. CQ is supported by National Institute of Health New Innovator Award DP2OD008371.

## REFERENCES

Alon U., 2003 Biological networks: the tinkerer as an engineer. Science 301: 1866–7.

Alonso J. M., Stepanova A. N., Leisse T. J., Kim C. J., Chen H., Shinn P., Stevenson D. K., Zimmerman J., Barajas P., Cheuk R., Gadrinab C., Heller C., Jeske A., Koesema E., Meyers C. C., Parker H., Prednis L., Ansari Y., Choy N., Deen H., Geralt M., Hazari N., Hom E., Karnes M., Mulholland C., Ndubaku R., Schmidt I., Guzman P., Aguilar-Henonin L., Schmid M., Weigel D., Carter D. E., Marchand T., Risseeuw E., Brogden D., Zeko A., Crosby W. L., Berry C. C., Ecker J. R., 2003 Genome-wide insertional mutagenesis of Arabidopsis thaliana. Science 301: 653–7.

Alonso-Blanco C., Andrade J., Becker C., Bemm F., Bergelson J., Borgwardt K. M., Cao J., Chae E., Dezwaan T. M., Ding W., Ecker J. R., Exposito-Alonso M., Farlow A., Fitz J., Gan X., Grimm D. G., Hancock A. M., Henz S. R., Holm S., Horton M., Jarsulic M., Kerstetter R. A., Korte A., Korte P., Lanz C., Lee C.-R., Meng D., Michael T. P., Mott R., Muliyati N. W., Nägele T., Nagler M., Nizhynska V., Nordborg M., Novikova P. Y., Picó F. X., Platzer A., Rabanal F. A., Rodriguez A., Rowan B. A., Salomé P. A., Schmid K. J., Schmitz R. J., Seren Ü., Sperone F. G., Sudkamp M., Svardal H., Tanzer M. M., Todd D., Volchenboum S. L., Wang C., Wang G., Wang X., Weckwerth W., Weigel D., Zhou X., 2016 1,135 Genomes Reveal the Global Pattern of Polymorphism in Arabidopsis thaliana. Cell 0.

Anwer M. U., Boikoglou E., Herrero E., Hallstein M., Davis A. M., James G. V., Nagy F., Davis S. J., 2014 Natural variation reveals that intracellular distribution of ELF3 protein is associated with function in the circadian clock. eLife 3: e02206.

Broman K. W., Wu H., Sen S., Churchill G. A., 2003 R/qtl: QTL mapping in experimental crosses. Bioinformatics 19: 889–890.

Chow B. Y., Helfer A., Nusinow D. A., Kay S. A., 2012 ELF3 recruitment to the PRR9 promoter requires other Evening Complex members in the Arabidopsis circadian clock. Plant Signal. Behav. 7: 170–3.

Escher D., Bodmer-Glavas M., Barberis A., Schaffner W., 2000 Conservation of Glutamine-Rich Transactivation Function between Yeast and Humans. Mol. Cell. Biol. 20: 27742782.

Fondon J. W., Hammock E. A. D., Hannan A. J., King D. G., 2008 Simple sequence repeats: genetic modulators of brain function and behavior. Trends Neurosci. 31: 328–34.

Gan X., Stegle O., Behr J., Steffen J. G., Drewe P., Hildebrand K. L., Lyngsoe R., Schultheiss S. J., Osborne E. J., Sreedharan V. T., Kahles A., Bohnert R., Jean G., Derwent P., Kersey P., Belfield E. J., Harberd N. P., Kemen E., Toomajian C., Kover P. X., Clark R. M., Ratsch G., Mott R., 2011 Multiple reference genomes and transcriptomes for Arabidopsis thaliana. Nature 477: 419–23.

Gemayel R., Vinces M. D., Legendre M., Verstrepen K. J., 2010 Variable tandem repeatsaccelerate evolution of coding and regulatory sequences. Annu. Rev. Genet. 44: 44577.

Hannan A. J., 2010 Tandem repeat polymorphisms: modulators of disease susceptibility and candidates for “missing heritability”. Trends Genet. 26: 59–65.

Herrero E., Kolmos E., Bujdoso N., Yuan Y., Wang M., Berns M. C., Uhlworm H., Coupland G., Saini R., Jaskolski M., Webb A., Gonçalves J., Davis S. J., 2012 EARLY FLOWERING4 recruitment of EARLY FLOWERING3 in the nucleus sustains the Arabidopsis circadian clock. Plant Cell 24: 428–43.

Hicks K. A., Millar A. J., Carre I. A., Somers D. E., Straume M., Meeks-Wagner D. R., Kay S. A., 1996 Conditional Circadian Dysfunction of the Arabidopsis early-flowering 3 Mutant. Science 274: 790–792.

Jacob F., 1977 Evolution and Tinkering. Science 196: 1161–1166.

Jain R., Rivera M. C., Lake J. a, 1999 Horizontal gene transfer among genomes: the complexity hypothesis. Proc. Natl. Acad. Sci. U. S. A. 96: 3801–6.

Jimenez-Gomez J. M., Corwin J. a., Joseph B., Maloof J. N., Kliebenstein D. J., 2011 Genomic Analysis of QTLs and Genes Altering Natural Variation in Stochastic Noise (G Gibson, Ed.). PLoS Genet. 7: e1002295.

Kashi Y., King D., Soller M., 1997 Simple sequence repeats as a source of quantitative genetic variation. Trends Genet. 13: 74–78.

Khattak A. K., 2014 Natural Variation in Arabidopsis thaliana Growth in Response to Ambient Temperatures: PhD Thesis.

Kleinboelting N., Huep G., Kloetgen A., Viehoever P., Weisshaar B., 2012 GABI-Kat SimpleSearch: new features of the Arabidopsis thaliana T-DNA mutant database. Nucleic Acids Res. 40: D1211–5.

Kolmos E., Herrero E., Bujdoso N., Millar A. J., Tóth R., Gyula P., Nagy F., Davis S. J., 2011 A Reduced-Function Allele Reveals That EARLY FLOWERING3 Repressive Action on the Circadian Clock Is Modulated by Phytochrome Signals in Arabidopsis. Plant Cell Online 23: 3230–3246.

Kover P. X., Valdar W., Trakalo J., Scarcelli N., Ehrenreich I. M., Purugganan M. D., Durrant C., Mott R., 2009 A Multiparent Advanced Generation Inter-Cross to Fine-Map Quantitative Traits in <italic>Arabidopsis thaliana</italic>. PLoS Genet 5: e1000551.

Lachowiec J., Queitsch C., Kliebenstein D. J., 2015 Molecular mechanisms governing differential robustness of development and environmental responses in plants. Ann. Bot.: mcv151.

Lander E. S., Botstein D., 1989 Mapping Mendelian Factors Underlying Quantitative Traits Using RFLP Linkage Maps. Genetics 121: 185–199.

Lempe J., Lachowiec J., Sullivan A. M., Queitsch C., 2013 Molecular mechanisms of robustness in plants. Curr. Opin. Plant Biol. 16: 62–9.

Li H., Handsaker B., Wysoker A., Fennell T., Ruan J., Homer N., Marth G., Abecasis G., Durbin R., 2009 The Sequence Alignment/Map format and SAMtools. Bioinforma. Oxf. Engl. 25: 2078–9.

Li H., 2013 *Aligning sequence reads, clone sequences and assembly contigs with BWA-MEM*. http://ArXiv:3.

Liu X. L., Covington M. F., Fankhauser C., Chory J., Wagner D. R., 2001 ELF3 encodes a circadian clock-regulated nuclear protein that functions in an Arabidopsis PHYB signal transduction pathway. Plant Cell 13: 1293–304.

Lu X., Murphy R. M., 2015 Asparagine Repeat Peptides: Aggregation Kinetics and Comparison with Glutamine Repeats. Biochemistry (Mosc.) 54: 4784–94.

Möckli N., Auerbach D., 2004 Quantitative β-galactosidase assay suitable for high-throughput applications in the yeast two-hybrid system. BioTechniques 36: 872876.

Nieto C., López-Salmerón V., Davière J.-M., Prat S., 2014 ELF3-PIF4 Interaction Regulates Plant Growth Independently of the Evening Complex. Curr. Biol. 25: 187–193.

Nusinow D. A., Helfer A., Hamilton E. E., King J. J., Imaizumi T., Schultz T. F., Farré E. M., Kay S. A., 2011 The ELF4-ELF3-LUX complex links the circadian clock to diurnal control of hypocotyl growth. Nature 475: 398–402.

Orr H. T., Zoghbi H. Y., 2007 Trinucleotide Repeat Disorders. Annu. Rev. Neurosci. 30: 575621.

Perutz M. F., Johnson T., Suzuki M., Finch J. T., 1994 Glutamine repeats as polar zippers: their possible role in inherited neurodegenerative diseases. Proc. Natl. Acad. Sci. 91: 5355–5358.

Press M. O., Carlson K. D., Queitsch C., 2014 The overdue promise of short tandem repeat variation for heritability. Trends Genet. 30: 504–512.

Press M. O., Lanctot A., Queitsch C., 2016 ELF3 polyQ variation in Arabidopsis thaliana reveals a PIF4-independent role in thermoresponsive flowering. bioRxiv: 038257.

Queitsch C., Carlson K. D., Girirajan S., 2012 Lessons from model organisms: phenotypic robustness and missing heritability in complex disease. (SM Rosenberg, Ed.). PLoS Genet. 8: e1003041.

R Core Team, 2016 R: A Language and Environment for Statistical Computing. R Foundation for Statistical Computing, Vienna, Austria.

Rowan B. A., Patel V., Weigel D., Schneeberger K., 2015 Rapid and inexpensive whole-genome genotyping-by-sequencing for crossover localization and fine-scale genetic mapping. G3 Bethesda Md 5: 385–98.

Rutherford S. L., Lindquist S., 1998 Hsp90 as a capacitor for morphological evolution. Nature 396: 336–42.

Sangster T. a, Bahrami A., Wilczek A., Watanabe E., Schellenberg K., McLellan C., Kelley A., Kong S. W., Queitsch C., Lindquist S., 2007 Phenotypic diversity and altered environmental plasticity in Arabidopsis thaliana with reduced Hsp90 levels. PloS One 2: e648.

Sangster T. a, Salathia N., Lee H. N., Watanabe E., Schellenberg K., Morneau K., Wang H., Undurraga S., Queitsch C., Lindquist S., 2008a HSP90-buffered genetic variation is common in Arabidopsis thaliana. Proc. Natl. Acad. Sci. U. S. A. 105: 2969–74.

Sangster T. a, Salathia N., Undurraga S., Milo R., Schellenberg K., Lindquist S., Queitsch C., 2008b HSP90 affects the expression of genetic variation and developmental stability in quantitative traits. Proc. Natl. Acad. Sci. U. S. A. 105: 2963–8.

Schaefer M. H., Wanker E. E., Andrade-Navarro M. A., 2012 Evolution and function of CAG/polyglutamine repeats in protein-protein interaction networks. Nucleic Acids Res. 40: 4273–87.

Sharma D., Sharma S., Pasha S., Brahmachari S. K., 1999 Peptide models for inherited neurodegenerative disorders: conformation and aggregation properties of long polyglutamine peptides with and withoutk interruptions. FEBS Lett. 456: 181–185.

Sievers F., Wilm A., Dineen D., Gibson T. J., Karplus K., Li W., Lopez R., McWilliam H., Remmert M., Söding J., Thompson J. D., Higgins D. G., 2011 Fast, scalable generation of high-quality protein multiple sequence alignments using Clustal Omega. Mol. Syst. Biol. 7: 539.

Stott K., Blackburn J. M., Butler P. J., Perutz M., 1995 Incorporation of glutamine repeats makes protein oligomerize: implications for neurodegenerative diseases. Proc. Natl. Acad. Sci. U. S. A. 92: 6509–13.

Sureshkumar S., Todesco M., Schneeberger K., Harilal R., Balasubramanian S., Weigel D., 2009 A genetic defect caused by a triplet repeat expansion in Arabidopsis thaliana. Science 323: 1060–3.

Szamecz B., Boross G., Kalapis D., Kovács K., Fekete G., Farkas Z., Lázár V., Hrtyan M., Kemmeren P., Groot Koerkamp M. J. A., Rutkai E., Holstege F. C. P., Papp B., Pál C., 2014 The Genomic Landscape of Compensatory Evolution. PLoS Biol. 12: e1001935.

Taipale M., Jarosz D. F., Lindquist S., 2010 HSP90 at the hub of protein homeostasis: emerging mechanistic insights. Nat. Rev. Mol. Cell Biol. 11: 515–28.

Tajima T., Oda A., Nakagawa M., Kamada H., Mizoguchi T., 2007 Natural variation of polyglutamine repeats of a circadian clock gene ELF3 in Arabidopsis. Plant Biotechnol. 24: 237–240.

Undurraga S. F., Press M. O., Legendre M., Bujdoso N., Bale J., Wang H., Davis S. J., Verstrepen K. J., Queitsch C., 2012 Background-dependent effects of polyglutamine variation in the Arabidopsis thaliana gene ELF3. Proc. Natl. Acad. Sci. U. S. A. 109: 19363–19367.

Usdin K., 2008 The biological effects of simple tandem repeats: Lessons from the repeat expansion diseases. Genome Res. 18: 1011–1019.

Wang Y., Lu J., Yu J., Gibbs R. A., Yu F., 2013 An integrative variant analysis pipeline for accurate genotype/haplotype inference in population NGS data. Genome Res. 23: 833–42.

Yoshida R., Fekih R., Fujiwara S., Oda A., Miyata K., Tomozoe Y., Nakagawa M., Niinuma K., Hayashi K., Ezura H., Coupland G., Mizoguchi T., 2009 Possible role of early flowering 3 (ELF3) in clock-dependent floral regulation by short vegetative phase (SVP) in Arabidopsis thaliana. New Phytol. 182: 838–50.

Yu J.-W., Rubio V., Lee N.-Y., Bai S., Lee S.-Y., Kim S.-S., Liu L., Zhang Y., Irigoyen M. L., Sullivan J. A., Zhang Y., Lee I., Xie Q., Paek N.-C., Deng X. W., 2008 COP1 and ELF3 control circadian function and photoperiodic flowering by regulating GI stability. Mol. Cell 32: 617–30.

Zhao L., Nakazawa M., Takase T., Manabe K., Kobayashi M., Seki M., Shinozaki K., Matsui M., 2004 Overexpression of LSH1, a member of an uncharacterised gene family, causes enhanced light regulation of seedling development. Plant J. 37: 694–706.

Zhao R., Davey M., Hsu Y.-C., Kaplanek P., Tong A., Parsons A. B., Krogan N., Cagney G., Mai D., Greenblatt J., Boone C., Emili A., Houry W. A., 2005 Navigating the chaperone network: an integrative map of physical and genetic interactions mediated by the hsp90 chaperone. Cell 120: 715–27.

